# Prediction of druggable proteins using machine learning and functional enrichment analysis: a focus on cancer-related proteins and RNA-binding proteins

**DOI:** 10.1101/825513

**Authors:** Andrés López-Cortés, Alejandro Cabrera-Andrade, Carlos M. Cruz-Segundo, Julian Dorado, Alejandro Pazos, Humberto Gonzáles-Díaz, César Paz-y-Miño, Yunierkis Pérez-Castillo, Eduardo Tejera, Cristian R. Munteanu

**Affiliations:** Centro de Investigación Genética y Genómica, Facultad de Ciencias de la Salud Eugenio Espejo, Universidad UTE, Mariscal Sucre Avenue, Quito 170129, Ecuador; RNASA-IMEDIR, Computer Science Faculty, University of Coruna, Coruna 15071, Spain; Red Latinoamericana de Implementación y Validación de Guías Clínicas Farmacogenómicas (RELIVAF-CYTED); Grupo de Bio-Quimioinformática, Universidad de Las Américas, Avenue de los Granados, Quito 170125, Ecuador; Carrera de Enfermería, Facultad de Ciencias de la Salud, Universidad de Las Américas, Avenue de los Granados, Quito 170125, Ecuador; Tecnológico de Estudios Superiores de Jocotitlán, Carretera Toluca-Atlacomulco KM 44.8 Ejido de San Juan y San Agustin, 50700 Jocotitlán, México; Centro de Investigación en Tecnologías de la Información y las Comunicaciones (CITIC), Campus de Elviña s/n 15071 A Coruña, Spain; Biomedical Research Institute of A Coruña (INIBIC), University Hospital Complex of A Coruña (CHUAC), 15006, A Coruña, Spain; Department of Organic Chemistry II, University of the Basque Country UPV/EHU, Leioa 48940, Biscay, Spain; IKERBASQUE, Basque Foundation for Science, Bilbao 48011, Biscay, Spain; Escuela de Ciencias Físicas y Matemáticas, Universidad de Las Américas, Avenue de los Granados, Quito 170125, Ecuador; Facultad de Ingeniería y Ciencias Agropecuarias, Universidad de Las Américas, Avenue de los Granados, Quito 170125, Ecuador

**Author notes:** These authors contributed equally to the study (Andrés López-Cortés, Alejandro Cabrera-Andrade). E-mail of authors: Andrés López-Cortés, Alejandro Cabrera-Andrade, Carlos M. Cruz-Segundo, Julian Dorado, Alejandro Pazos, Humberto Gonzáles-Díaz, César Paz-y-Miño, Yunierkis Pérez-Castillo, Eduardo Tejera, Cristian R. Munteanu. Correspondence: Andrés López-Cortés, MSc., Centro de Investigación Genética y Genómica, Facultad de Ciencias de la Salud Eugenio Espejo, Universidad UTE, Mariscal Sucre Avenue, Quito 170129, Ecuador.

**Keywords:** breast cancer, cancer-driving proteins, clinical trials, druggable proteins, machine learning, RNA-binding proteins, support vector machine

## Abstract

**Background:** Druggable proteins are a trending topic in drug design. The druggable proteome can be defined as the percentage of proteins that have the capacity to bind an antibody or small molecule with adequate chemical properties and affinity. The screening and *in silico* modeling are critical activities for the reduction of experimental costs.

**Methods:** The current work proposes a unique prediction model for druggable proteins using amino acid composition descriptors of protein sequences and 13 machine learning linear and non-linear classifiers. After feature selection, the best classifier was obtained using the support vector machine method and 200 tri-amino acid composition descriptors.

**Results:** The high performance of the model is determined by an area under the receiver operating characteristics (AUROC) of 0.975 ± 0.003 and accuracy of 0.929 ± 0.006 (3-fold cross-validation). Regarding the prediction of cancer-associated proteins using this model, the best ranked druggable predicted proteins in the breast cancer protein set were CDK4, AP1S1, POLE, HMMR, RPL5, PALB2, TIMP1, RPL22, NFKB1 and TOP2A; in the cancer-driving protein set were TLL2, FAM47C, SAGE1, HTR1E, MACC1, ZFR2, VMA21, DUSP9, CTNNA3 and GABRG1; and in the RNA-binding protein set were PLA2G1B, CPEB2, NOL6, LRRC47, CTTN, CORO1A, SCAF11, KCTD12, DDX43 and TMPO.

**Conclusions:** This powerful model predicts several druggable proteins which should be deeply studied to find better therapeutic targets and thus improve clinical trials. The scripts are freely available at https://github.com/muntisa/machine-learning-for-druggable-proteins.

## INTRODUCTION

The human genome is made of ∼20,000 protein-coding genes, but not all proteins are suitable drug targets [1–3]. The druggable proteome can be defined as the percentage of proteins that have the capacity to bind an antibody or small molecule with adequate chemical properties and affinity [4]. Druggability is the property of a druggable molecule (i.e., a biological target) by virtue of which it elicits a favorable clinical response when it contacts a drug-like compound [5]. It is estimated that 60% of the small molecule drug discovery projects fail during the hit-to-lead phase because the biological target is found to be not druggable [5, 6]. Therefore, it is important to be able to predict how druggable a novel target is in early drug discovery because only 10% of the human genome represents druggable targets, and only half of those are relevant to diseases [7].

According to Gashaw *et al*., an ideal drug target should have the following properties: freedom to operate (i.e., lack of competitive binding), the existence of a biomarker to monitor its efficacy, differential expression across the body for specific targeting, low impact on the modulation of physiological conditions, capacity to modify a disease, and favorable assayability for high throughput screening [7, 8].

Regarding all the protein-coding genes in the human genome, approximately 3,000 are estimated to be part of the druggable genome. Yet, drugs approved by the US Food & Drug Administration (FDA) target just twenty percent of the druggable proteins [9]. FDA has approved the use of 672 (100%) drugs classified by their protein class in enzymes (260; 39%), transporters (149; 22%), G-protein coupled receptors (98; 15%), CD markers (71; 11%), voltage-gated ion channels (49; 7%), nuclear receptors (24; 4%), among others [10, 11]. Drugs that inactivate the protein target are known as antagonists, while drugs that activate the protein target are called agonists. Regarding the cellular location of targets for FDA approved drugs based on a variety of transmembrane and signal protein prediction methods, 250 (37%) were integral membrane (IM), 201 (30%) were intracellular, 101 (15%) were single pass transmembrane (SPTM), 83 (12%) were secreted, 28 (4%) were membrane and secreted isoforms, and 9 (1%) were IM/SPTM [4, 10, 11].

The low number of approved drugs up to date is due to several factors such as the complexity in the experimentation of all proteins and fragments of nucleic acids, and the lack of knowledge of various diseases at molecular level. Due to these facts, computational models that can predict drug targets with high sensitivity while maintaining a high specificity on a genome-wide scale would be highly welcomed [5]. Additionally, it is possible to combine big data such as metabolic and gene regulatory networks, protein-protein interactions, multi-omics information and gene expression profiles with data mining tools like machine learning (ML) to build predictive models in order to identify biological relevant patterns that confer druggability to potential drug targets [12].

Classification models have been published for the prediction of protein activities such as anti-angiogenic [13], anti-cancer [14], enzyme class [15], epitope [16], signaling [17], lectins [18], anti-oxidant [19], and druggability [12, 20–26]. Therefore, the aim of this study was to build an effective machine learning classifier to predict druggability of breast cancer (BC) proteins, cancer-driving proteins and RNA-binding proteins (RBPs).

## METHODS

In Fig. 1, the general flow chart of the proposed methodology is presented. In the first step, a database with druggable and non-druggable proteins was constructed. Next, three families of protein composition descriptors were calculated using Rcpi (R package) [27]: 20 amino acid composition (AC), 400 di-amino acid composition (DC) and 8000 tri-amino acid composition (TC). In the next step, jupyter notebooks based on python/sklearn [28] built 13 types of ML classifiers by combining the 3 families of descriptors (AC, DT, TC) with five different feature selection methods and with different parameters. The classifiers were Gaussian Naive Bayes (GNB) [29], k-nearest neighbors algorithm (KNeighborsClassifier) [30], linear discriminant analysis (LinearDiscriminantAnalysis) [31], support vector machine (linear and non-linear based on radial basis functions (RBF))[32], logistics regression (LogisticRegression) [33], multilayer perceptron / neural network (MLPClassifier) with 20 neurons in one hidden layer [34], decision tree (DecisionTreeClassifier) [35], random forest (RandomForestClassifier) [36], XGBoost - an optimized distributed gradient boosting library (XGBClassifier) [37], Gradient Boosting for classification (GradientBoostingClassifier) [38], AdaBoost classifier (AdaBoostClassifier) [39], and Bagging classifier (BaggingClassifier) [40]. The future selection methods were principal component analysis (PCA) [41], feature selection according to a percentile of the highest scores (SelectPercentile) with f_classif (ANOVA F-value between label/feature for classification tasks), feature selector that removes features with variance lower than a cutoff (VarianceThreshold), feature selection using Linear Support Vector classification (LinearSVC) and extra-trees classifier (ExtraTreesClassifier).

**Fig. 1.**
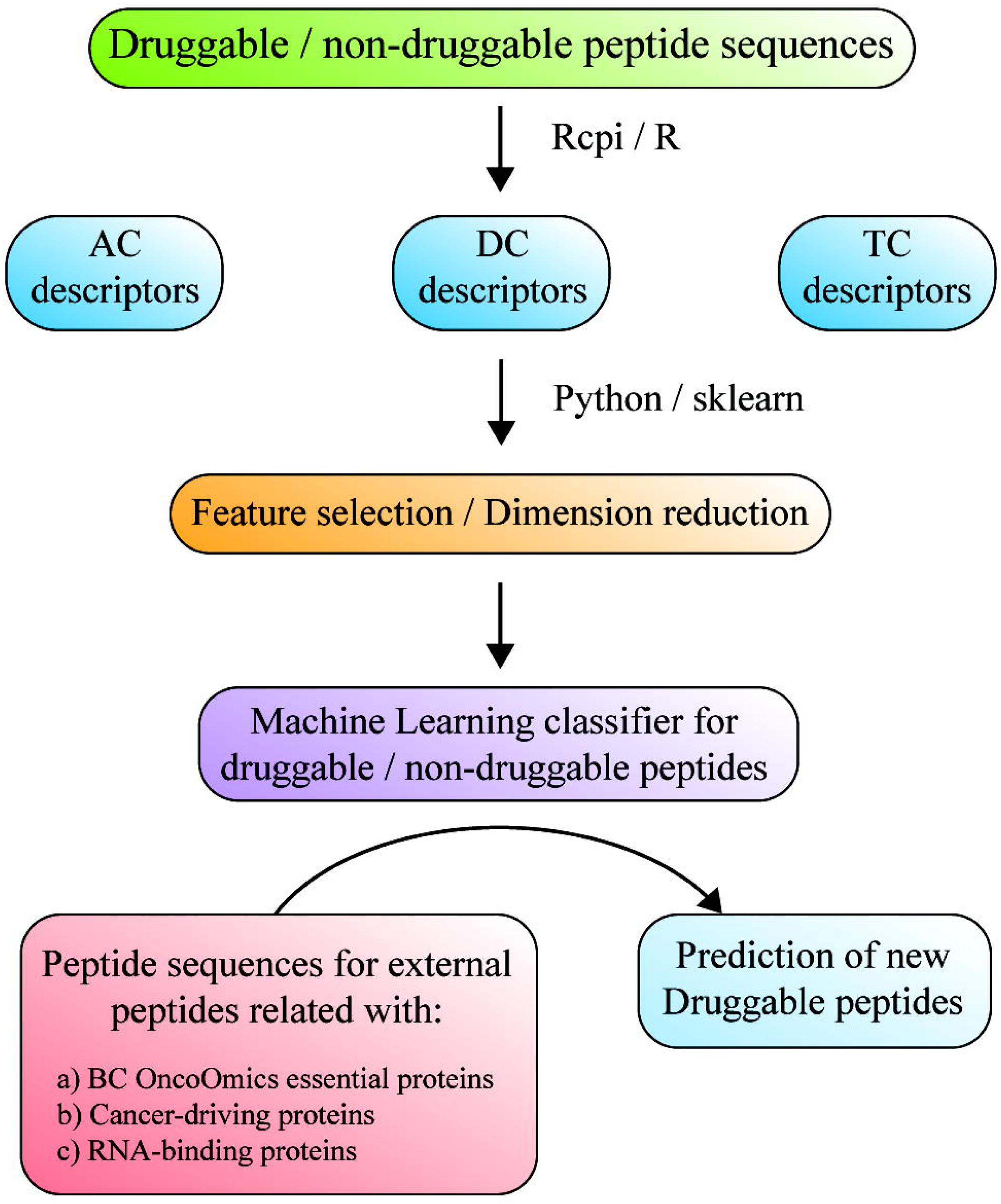
Flow chart of methodology for druggable protein prediction. AC, amino acid composition; DC, di-amino acid composition; TC, tri-amino acid composition; BC, breast cancer.

GNB is a probabilistic classifier based on the Bayes’ theorem that has a naive assumption: all the features are independent [29]. k-nearest neighbors algorithm (KNN) is a non-parametric classifier that assigns an unclassified sample to the same class as the nearest of k samples in the training set (this project uses k=3) [30]. Linear discriminant analysis (LDA) represents a linear classifier that fits the class conditional densities to input features by using Bayes’ rule [31]. Support vector machine (SVM linear and SVM) is non-linearly mapping input features to a higher dimensionality space, where a linear decision surface can be established [32]. The real problems are generally non-linear and SVM is solving this problem with the Gaussian radial basis (RBF) nonlinear kernel functions.

Logistics regression (LR) is a linear classifier that estimate the probability of a binary response by using different weights [33]. Multilayer perceptron (MLP) is a class of neural networks with artificial neurons, a hidden layer that is able to combine linear and nonlinear activation functions [34]. Decision tree (DT) (DecisionTreeClassifier) is a structure of decision rules inferred from the input features (classification rules = paths from root to leaf) [35]. Random forest (RF) is an ensemble method by combining parallel decision trees and it offers low-bias, low correlation between individual trees, and high variance [36].

XGBoost (XGB) is another type of ensemble method but it is using sequential weak trees to correct the classification errors [37]. Gradient Boosting for classification (GB) is a classical boot method based on the same sequential weak classifiers [38]. AdaBoost classifier (AdaB) is a meta-estimator that starts the fitting with a classifier based on the original dataset and then adds additional copies of the original classifier with adjusted weights for the incorrectly classified instances [39]. Bagging classifier (Bagging) is similar with AdaB but the additional classifiers are based on subsets of the original dataset [40].

The machine learning prediction model was constructed from two protein sets, the positive set was made up of 666 druggable proteins with FDA-approved drugs according to the DrugBank database (www.drugbank.ca) [10, 11, 42]. On the other hand, the negative protein set was made up of 219 non-druggable proteins [43]. Tables S1 and S2 detail the gene symbol and gene ID of all druggable and non-druggable proteins, and Tables S3 and S4 detail the FASTA sequences of all druggable and non-druggable proteins analyzed in this study.

Three lists of cancer-related proteins were scanned with the final ML prediction model: 230 were BC essential proteins [44], 2353 cancer-driving proteins were taken from the Network of Cancer Genes [45], and 1365 were RBPs [45, 46] (Tables S5-S7).

After the calculation of amino acid composition descriptors, the datasets contained 885 proteins (666 positive set, 219 negative set). The druggable class was labeled with 1, and the non-druggable class with 0. Due to the unbalanced datasets, a SMOTE filter was used [47]. A 3-fold cross-validation (CV) method was used to build the ML classifiers. A pipeline was constructed for each fold: a) scaler: the training set was scaled with StandardScaler and the test set was transformed to the same scale; b) feature selection / feature dimension reduction: the training set dimensionality was reduced by using or not a feature selection method (i.e., LinearSVC) or a feature dimension reduction (i.e., PCA); c) outer-CV: cross_val_score has been used to calculate the AUROC scores for the 13 ML methods in all splits; d) mean values and standard deviation (SD) of area under the receiver operating characteristics (AUROC) [48] for each ML classifier were printed (test subset).

The best model for predictions was chosen by taking into account several aspects: mean AUROC, SD of AUROC, number of features, and the type model features (original or transformed). The scripts are automatically calculating all the results and plotting the boxplots, and are freely available from https://github.com/muntisa/machine-learning-for-druggable-proteins.

After the screening of the three cancer-related protein sets through the ML model, the druggable predicted proteins were analyzed by using g:Profiler (https://biit.cs.ut.ee/gprofiler/) in order to obtain significant annotations (FDR < 0.001) related to gene ontology (GO) terms, pathways and disease phenotypes [49]. Finally, circos plots were created to show the current status of clinical trials of the best druggable predicted proteins according to the Open Targets Platform (https://www.targetvalidation.org). This platform is a comprehensive and robust data integration for access to and visualization of potential drug targets associated with cancer [50].

## RESULTS AND DISCUSSION

The project is proposing innovative classification models to predict new druggable proteins by using three families of protein composition descriptors calculated with Rcpi: AC, DC and TC. Jupyter notebooks based on python/sklearn were used to build classifiers using 13 types of ML classifiers and five types of feature selection methods, with different parameters (Fig. 1). The scripts used AUROC to quantify the classification performance. We tested models with 20, 100, 200, 400 and mixed numbers of features. The presented AUROC are mean values for 3-fold CV.

Fig. 2 presents AUROC values (test subset) for classifier using only 20 features: AC = AC descriptors without feature selection, DC-LinearSCV20 = DC descriptors obtained with LinearSCV feature selection, DC-PCAn20 = PCA features obtained from DC, TC-Percn20 = TC descriptors selected with SelectPercentile(f_classif, percentile=0.25), and TC-LinearSVC20 = TC descriptors selected with LinearSCV. We can observe that even with pure 20 AC descriptors (AC) and SVM it is possible to obtain an AUROC of 0.926. The best performance is obtained with SVM (RBF) and 20 PCA components from the 400 DC descriptors - AUROC = 0.958.

**Fig. 2.**
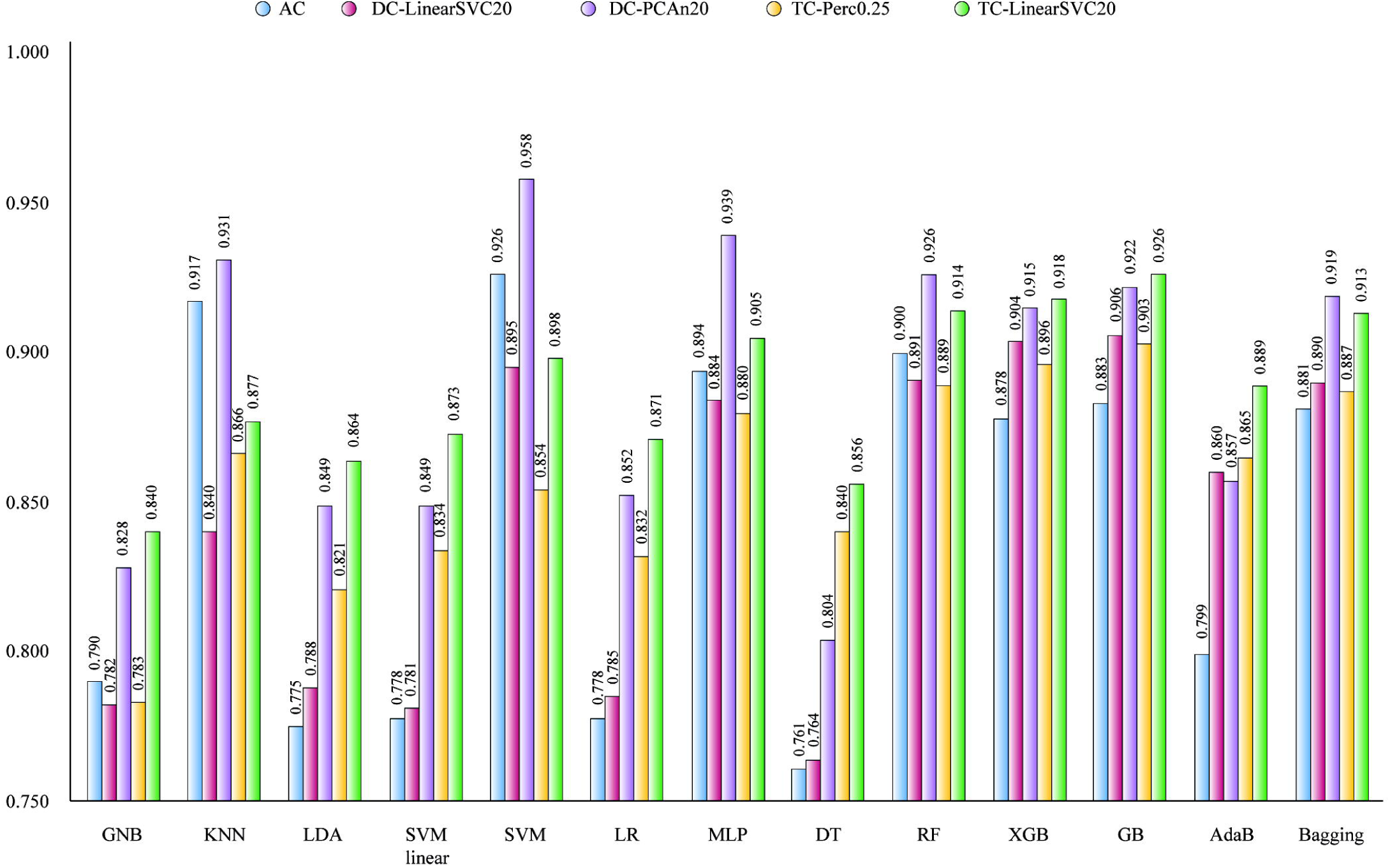
Mean AUROC of classifiers based on 20 input features (3-fold CV). GNB, Gaussian Naive Bayes; KNN, K-nearest neighbors algorithm; LDA, Linear discriminant analysis; SVM, Super vector machine; LR, Logistic regression; MLP, Multilayer perception; DT, Decision tree; RF, Random forest; XGB, XGBoost; GB, Gradient boosting; AdaB, AdaBoost classifier.

In Fig. 3 the number of inputs increased to 100: DC-PCAn100 = PCA transformed of 400 DC descriptors, TC-Perc1.25 = TC selected descriptors with SelectPercentile(f_classif, percentile=1.25), TC-LinearSVC100 = TC selected with LinearSVC, TC-PCA200LinearSVC100 = 100 selected features with LinearSVC from 200 PCA components of 8000 TC descriptors. By increasing the number of features to 100, the AUROC increases to 0.976 for the same SVM (RBF) and TC-PCA200LinearSVC100.

**Fig. 3.**
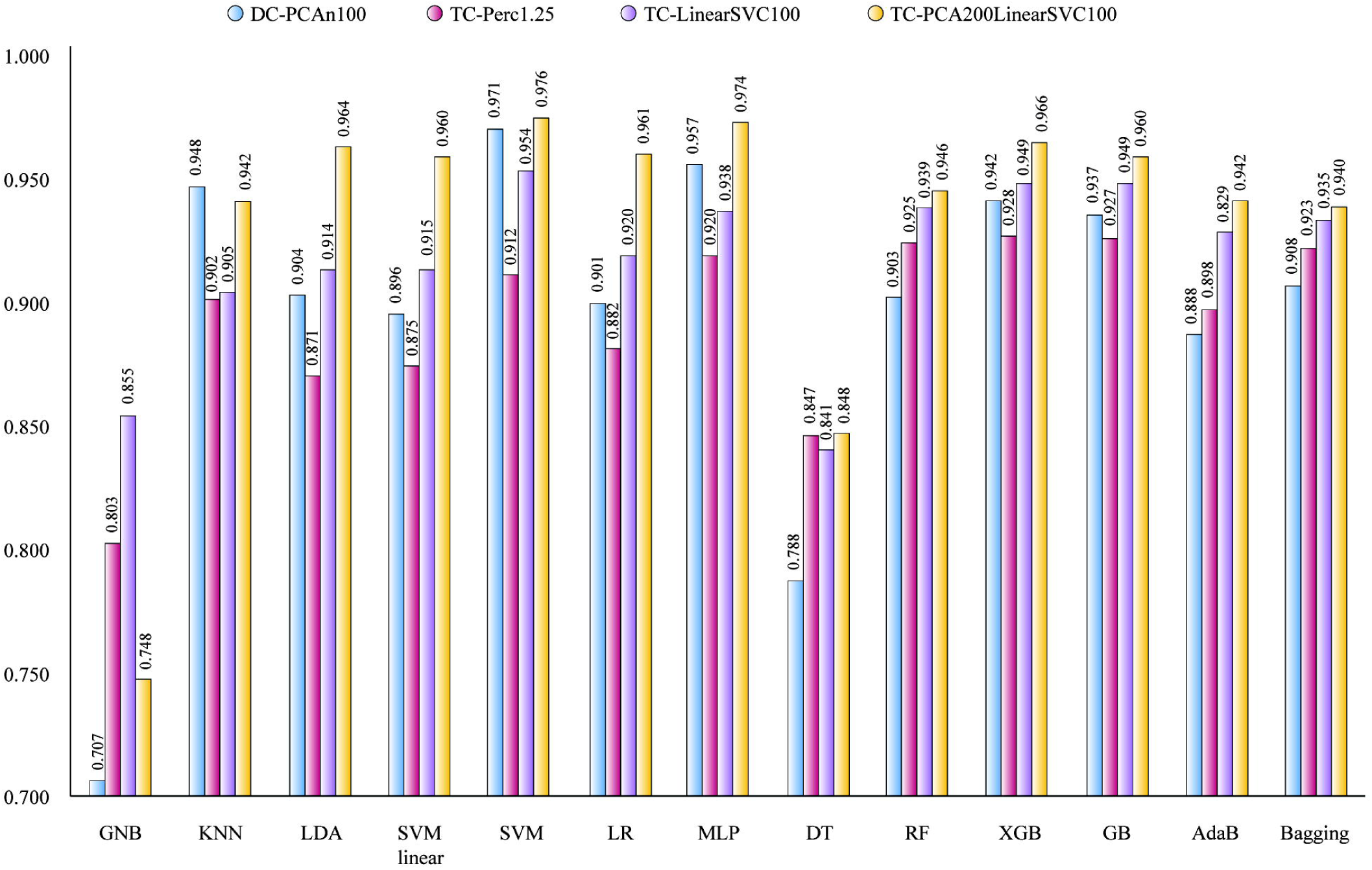
Mean AUROC of classifiers based on 100 input features (3-fold CV). GNB, Gaussian Naive Bayes; KNN, K-nearest neighbors algorithm; LDA, Linear discriminant analysis; SVM, Super vector machine; LR, Logistic regression; MLP, Multilayer perception; DT, Decision tree; RF, Random forest; XGB, XGBoost; GB, Gradient boosting; AdaB, AdaBoost classifier.

Fig. 4 presents the AUROC values for classifiers based on a double number of inputs (200 features): DC-PCAn200 = PCA transformed of 400 DC descriptors, DC-Perc50 = DC selected with SelectPercentile, TC-PCAn200 = 200 PCA components of 8000 transformed TC descriptors, TC-Perc2.5 = TC selected descriptors with SelectPercentile(f_classif, percentile=2.5), TC-LinearSVC200 = TC selected with LinearSVC. The same PCA mixed with SVM for DC-PCAn200 are resulting the best classifier with AUROC of 0.981. Additional results could be found in Table S8. If we will use all 400 DC descriptors with SVM, the mean AUROC could reach 0.982 ± 0.0021. Even more, with pure 8000 TC and SVM linear, the mean AUROC become 0.992 ± 0.0028. For a classification model, it should be avoided that the number of input features is more than the number of data instances. In addition, we tried to choose pure descriptors, not PCA transformed. Therefore, we made a compromise and we choose as the best model for the next predictions for proteins related to cancer: 200 TC descriptors selected with LinearSVC, non-linear SVM classifier with 0.975 ± 0.003 and accuracy of 0.929 ± 0.006 (3-fold cross-validation). The list of the 200 selected features is provided in the jupyter notebooks.

**Fig. 4.**
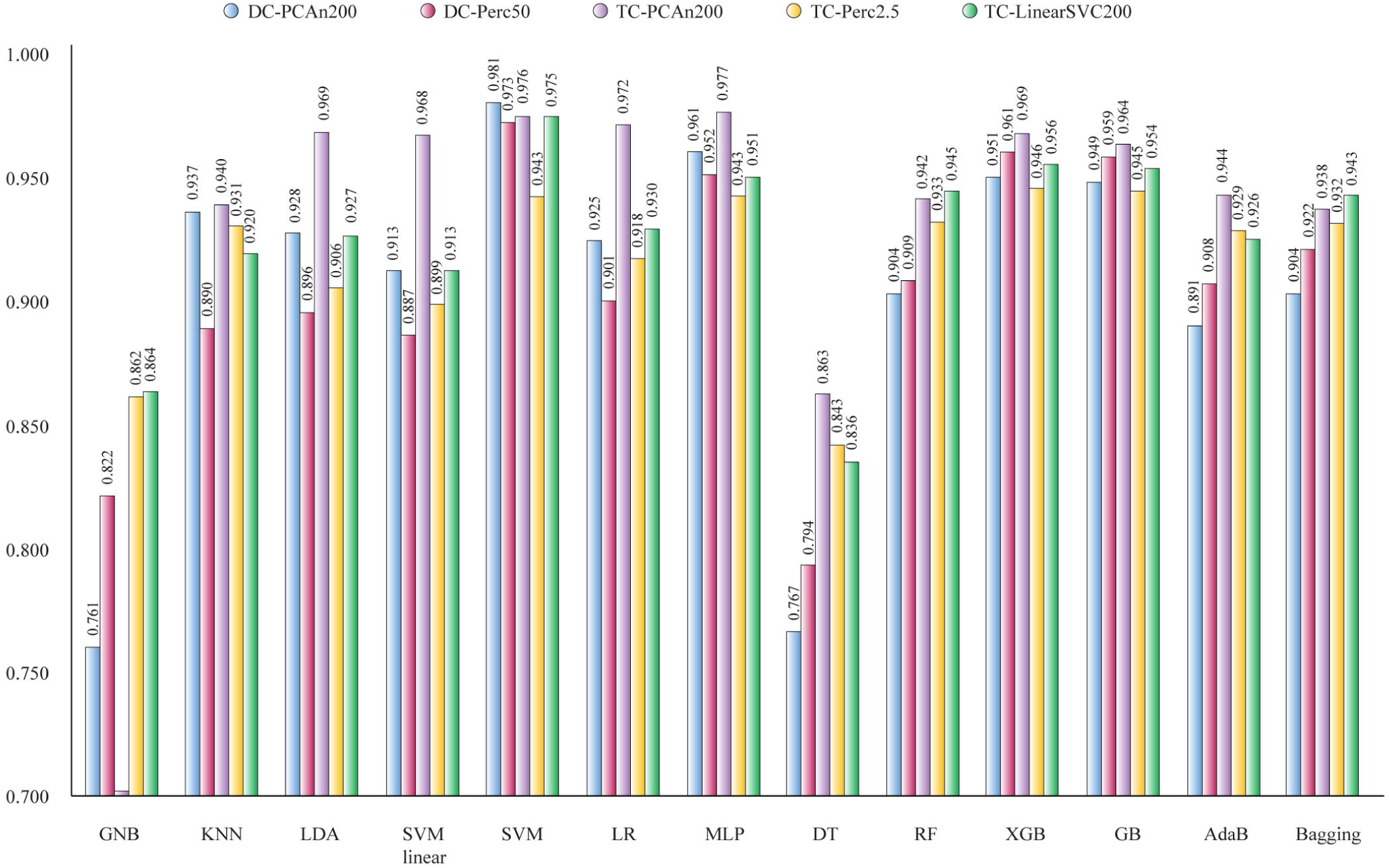
Mean AUROC of classifiers based on 200 input features (3-fold CV). GNB, Gaussian Naive Bayes; KNN, K-nearest neighbors algorithm; LDA, Linear discriminant analysis; SVM, Super vector machine; LR, Logistic regression; MLP, Multilayer perception; DT, Decision tree; RF, Random forest; XGB, XGBoost; GB, Gradient boosting; AdaB, AdaBoost classifier.

The three sets of proteins related to breast cancer proteins, cancer-driving proteins and RNA-binding proteins were evaluated for druggable features. Therefore, all these protein sequences have been transformed into scaled 200 selected TC descriptors. The results of these evaluations are provided in Tables S5-S7.

### Breast cancer essential proteins

After obtaining the list of proteins predicted as druggable, we performed a functional enrichment analysis through g:Profiler to interpret the gene ontology of these proteins [49]. Figs 5A and 5B show the enrichment map of the 209 BC proteins positively predicted as druggable. g:Profiler searches for a collection of proteins representing pathways, GO terms, and disease phenotypes [49]. The most significant false discovery rate (FDR < 0.001) GO: molecular functions were protein tyrosine kinase activity, protein binding, and enzyme binding; the most significant GO: biological processes were cell population proliferation, positive regulation of macromolecule metabolic process, and positive regulation of nitrogen compound metabolic process; the most significant Kyoto Encyclopedia of Genes and Genomes (KEGG) pathways were pathways in pathways in cancer, EGFR tyrosine kinase inhibitor resistance and MAPK signaling pathway [51, 52]; the most significant WikiPathways were pathway in glioblastoma, BC pathway and integrated BC pathway [53]; finally, the most significant Human Phenotype Ontology terms were cafe-au-late spot, neoplasm of the gastrointestinal tract, and breast carcinoma [54] (Table S9). The top 20 BC essential proteins predicted as druggable were CDK4, AP1S1, POLE, HMMR, RPL5, PALB2, TIMP1, RPL22, NFKB1, TOP2A, CTNNB1, BRCA2, TAF1, MLH1, NQO1, TOP3A, PIK3R1, RRM1, CDH1, and MMP2 (Fig. 5C). According to the Open Targets Platform [50], 14 of 20 best-ranked proteins do not have clinical trials in BC, TOP2A has 257, CDK4 has 125, POLE has 106, RRM1 has 105, PIK3R1 has 40, and MMP2 has 1 (Table S10, Figs. 5C and 5D). For instance, TOP2A has four clinical trials in phase 4 using doxorubicin, a small molecule that is inhibitor of the DNA topoisomerase II [50].

**Fig. 5.**
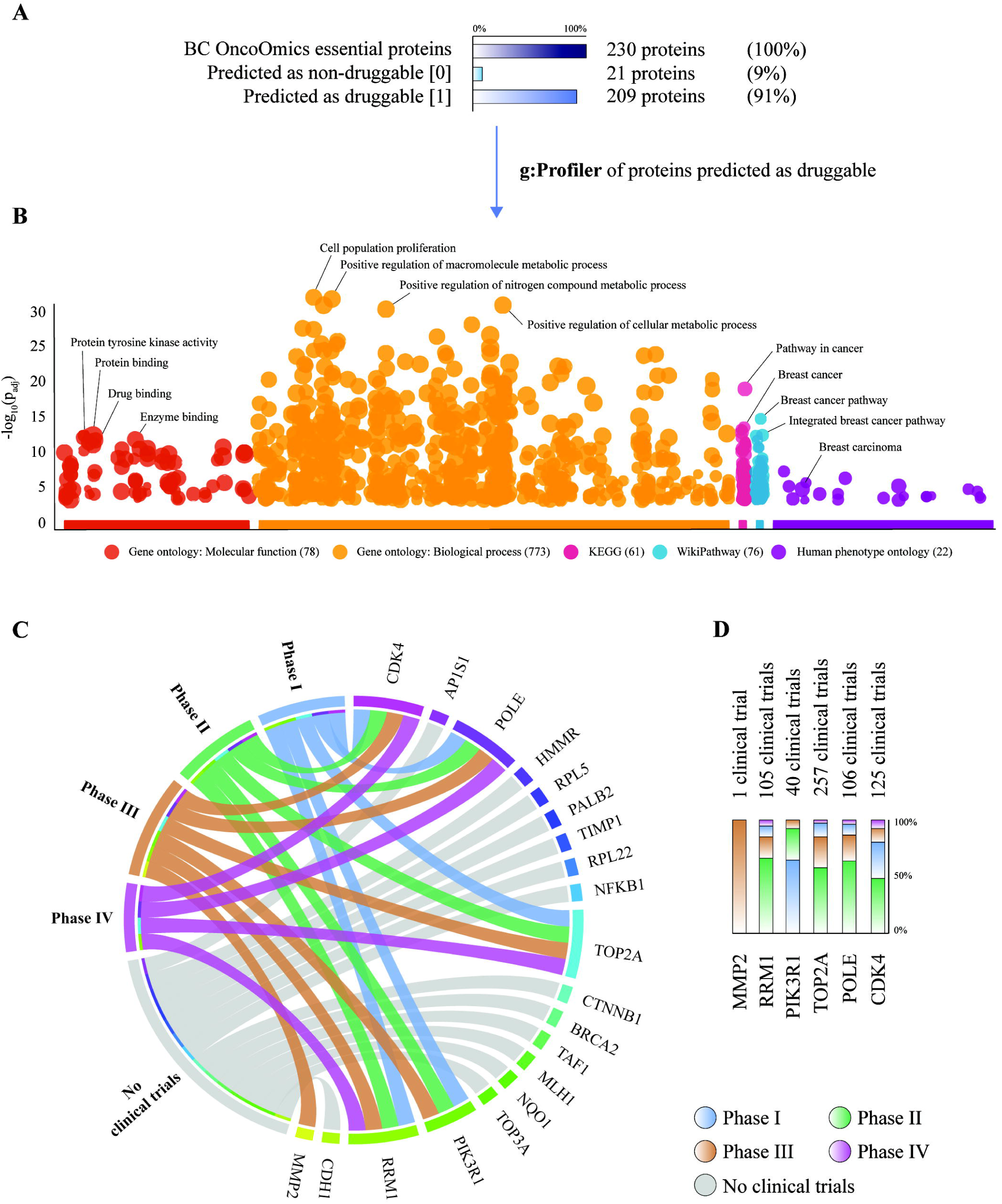
Breast cancer OncoOmics essential proteins. A) Percentage of proteins predicted as druggable and non-druggable. B) g:Profiler enrichment of BC proteins predicted as druggable. C) Circos plot of the 20 best-ranked BC proteins predicted as druggable, and its correlation with clinical trials. D) Phase and number of clinical trials per protein.

In our previous study [44], we revealed essential proteins associated with BC pathogenesis through deep analyses of genetic alterations, signaling pathways [52], protein-protein interaction networks [55, 56], protein expression [57, 58], dependency maps [57], and enrichment maps [49] of previously prioritized genes by the Consensus Strategy [59], the Pan-Cancer Atlas project [60–64], the Pharmacogenomics Knowledgebase (PharmGKB) [2], and the Cancer Genome Interpreter [65]. The druggable prediction of BC proteins through ML approaches will provide us relevant information regarding potential biomarkers, therapeutic targets and future clinical trials, avoiding ethnicity bias and improving cancer pharmacogenomics and precision medicine worldwide [66–76].

### Cancer-driving proteins

Figs. 6A and 6B show the enrichment map of the 2094 cancer-driving proteins positively predicted as druggable. Regarding g:Profiler [49], the most significant GO: molecular functions with a FDR < 0.001 were sequence-specific DNA binding, double-stranded DNA binding, and transcription regulatory region DNA binding; the most significant GO: biological processes were positive regulation of macromolecule metabolic process, positive regulation of nitrogen compound metabolic process, and positive regulation of metabolic process; the most significant KEGG pathways were EGFR tyrosine kinase inhibitor resistance, pathway in cancer, and transcriptional misregulation in cancer [51, 52]; additionally, the most significant term according to the Human Phenotype Ontology was somatic mutation [54] (Table S11). The top 20 cancer-driving proteins predicted as druggable were TLL2, FAM47C, SAGE1, HTR1E, MACC1, ZFR2, VMA21, DUSP9, CTNNA3, GABRG1, ZNF608, BIRC6, ARAF, ZNF483, TFRC, PCDHB6, DOCK4, TRPM3, TRAF6 and POF1B (Fig. 6C). Regarding the Open Targets Platform [50], 17 of 20 best-ranked proteins do not have clinical trials in all cancer types, GABRG1 has 34, TFRC has 10, and ARAF has 2 (Table S12, Figs. 6C and 6D). For instance, GABR1 has two clinical trials in phase 4 using propofol and sevoflurane, small molecules that belong to the GABA-A receptor class [50].

**Fig. 6.**
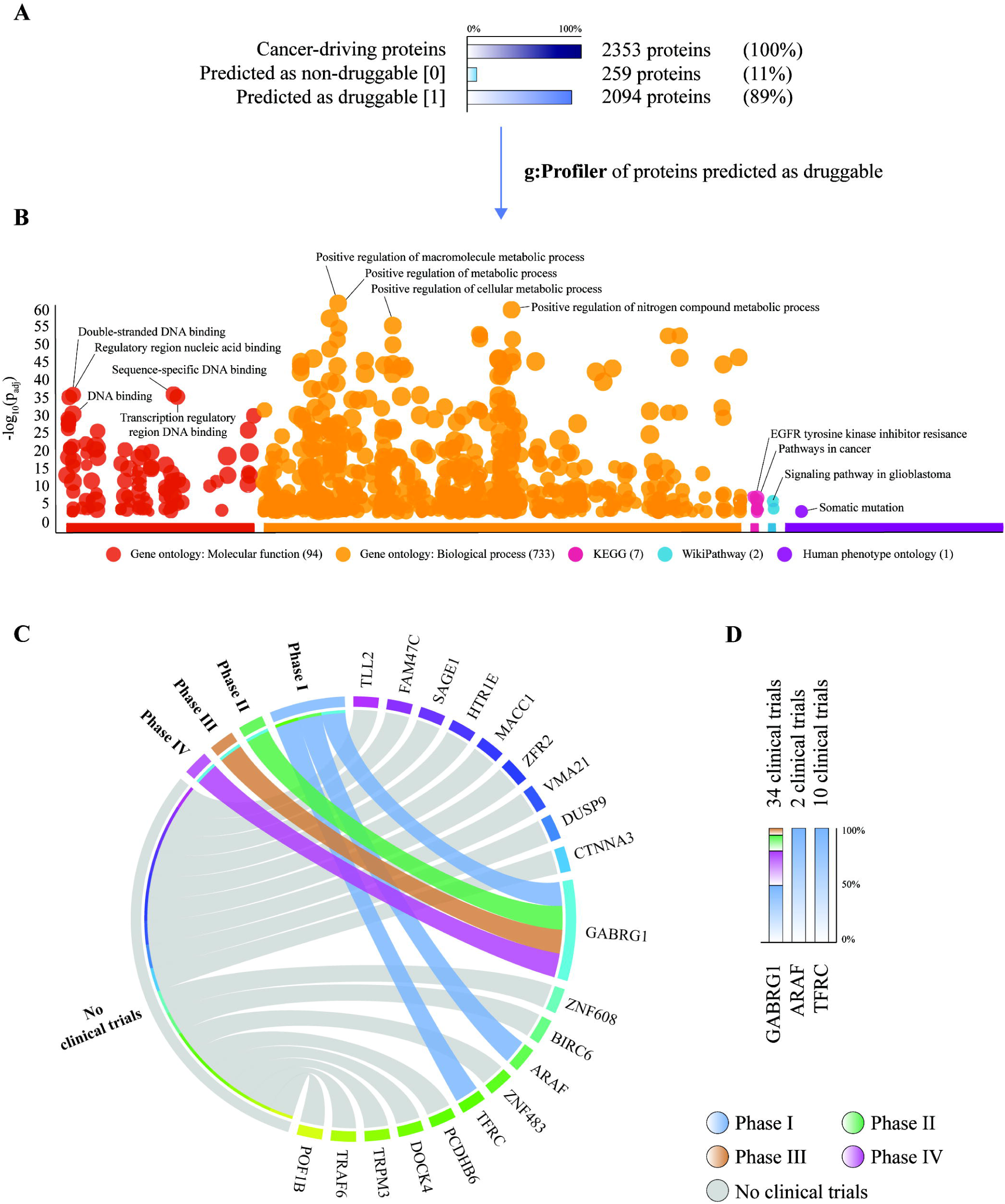
PanCancer-driving proteins. A) Percentage of proteins predicted as druggable and non-druggable. B) g:Profiler enrichment of PanCancer proteins predicted as druggable. C) Circos plot of the 20 best ranked PanCancer proteins predicted as druggable, and its correlation with clinical trials. D) Phase and number of clinical trials per protein.

The cancer-driving proteins were taken from The Network of Cancer Genes [45], a manually curated repository whose somatic modifications have known or predicted cancer driver roles. Fig. 7 shows the correlation between the best druggable predicted proteins and the cancer-driving proteins of 25 tissues of different primary sites (full list detailed in Table S13). The best druggable predicted protein per cancer type was RPL22 for adrenal gland cancer, ZFR2 for bladder cancer, TLL2 for blood cancer, CDKN2A for bone cancer, GABRG1 for brain cancer, PMS2 for breast cancer, KRAS for cervix cancer, GORASP1 for colorectal cancer, TRAF6 for esophagus cancer, FAT1 for head and neck cancer, ARAF for hepatobiliary cancer, CUL3 for kidney cancer, CTNNA3 for lung cancer, LDLRAD1 for melanoma, KRAS for ovary cancer, SAGE1 for pancreas cancer, FAT1 for penis cancer, CDKN2A for peripheral nervous system cancer, CDKN2A for prostate cancer, RSPH14 for small intestine cancer, HTR1E for stomach cancer, NDUFV2 for testis cancer, CDKN2A for thymus cancer, MACC1 for thyroid cancer, and USP9X for uterus cancer.

**Fig. 7.**
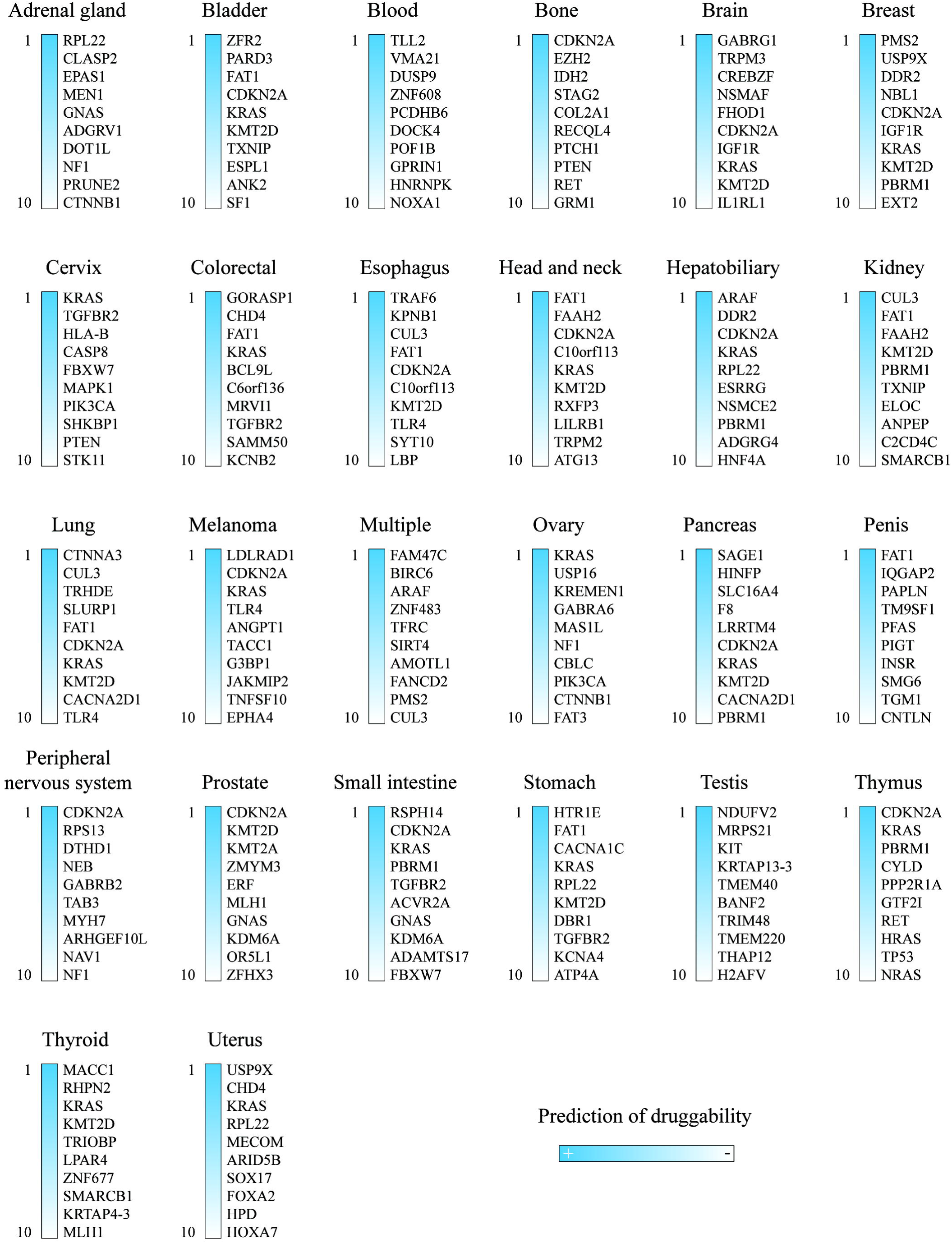
Top 10 druggable predicted proteins per cancer type.

### RNA-binding proteins

Figs. 8A and 8B show the enrichment map of the 1210 RBPs positively predicted as druggable. Regarding g:Profiler [49], the most significant (FDR < 0.001) GO: molecular functions were RNA binding, nucleic acid binding, and heterocyclic compound binding; the most significant GO: biological processes were RNA processing, ribonucleoprotein complex biogenesis, and mRNA metabolic process; the most significant KEGG pathways were spliceosome and ribosome biogenesis in eukaryotes [51, 52]; finally, the most significant WikiPathway was mRNA processing [53] (Table S14). The top 20 RBPs predicted as druggable were PLASG1B, CREB2, NOL6, LRRC47, CTTN, CORO1A, SCAF11, KCTD12, DDX43, TMPO, SARS2, SARNP, MEX3D, ZNF106, MTPAP, PRPF40A, YWHAZ, SEPTIN11, DDX46, and ANXA7 (Fig. 8C). According to the Open Targets Platform [50], the 20 best ranked proteins do not have clinical trials in cancer (Figs. 8C and 8D).

**Fig. 8.**
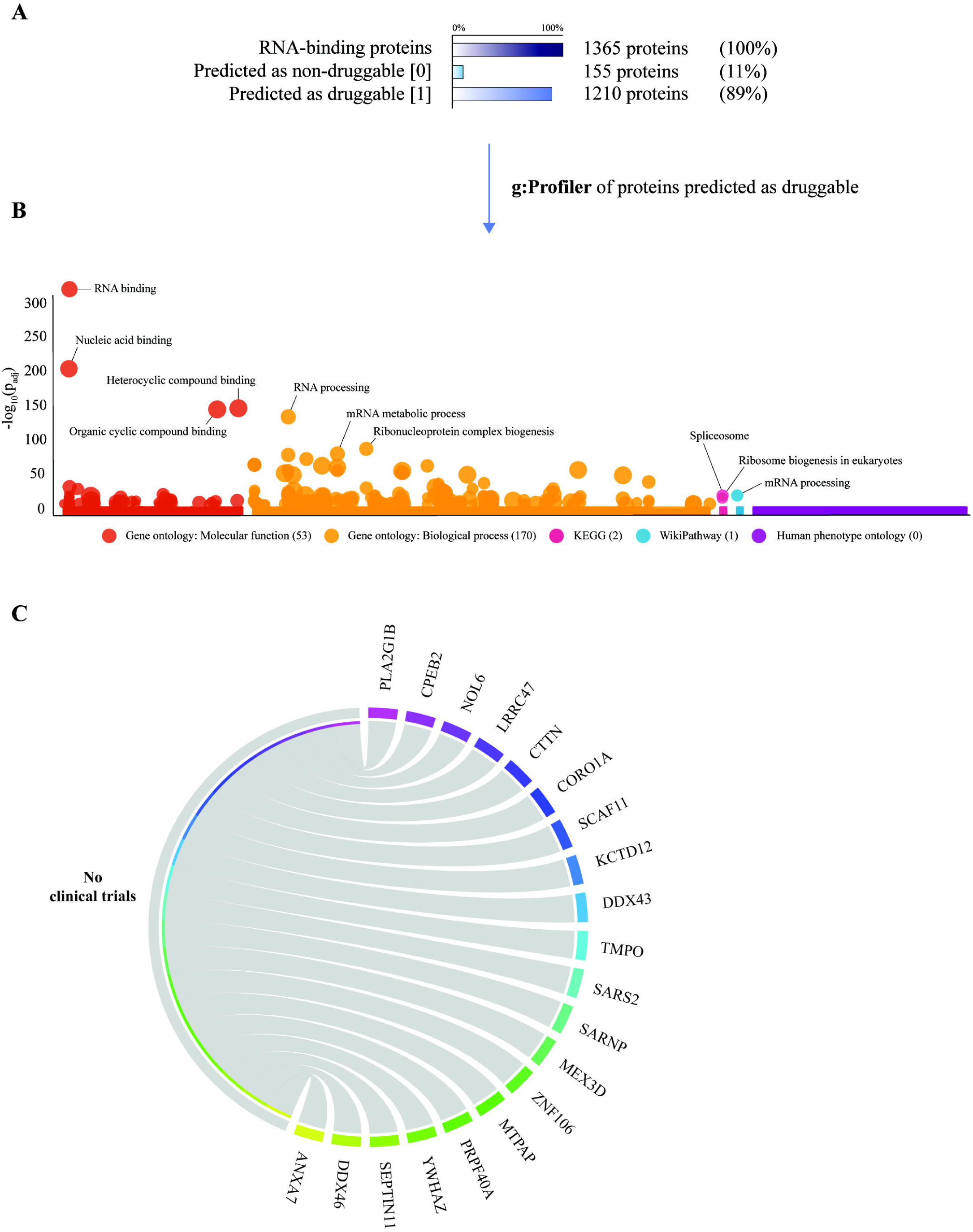
RNA-binding proteins. A) Percentage of proteins predicted as druggable and non-druggable. B) g:Profiler enrichment of RNA-binding proteins predicted as druggable. C) Circos plot of the 20 best ranked RNA-binding proteins predicted as druggable, and its correlation with clinical trials. D) Phase and number of clinical trials per protein.

RNA biology is an under-investigated field of cancer even though pleiotropic changes in the transcriptome are key feature of cancer cells [77]. RBPs are part of post-transcriptional RNA regulons. These RNA regulons are formed by ribonucleoproteins which interact with non-coding RNAs, metabolites, trans-elements and untranslated sequence elements found in mRNAs [78–81]. RBPs are able to control every aspect of RNA metabolism: degradation of mRNA, stability, polyadenylation, capping, translation, nucleocytoplasmic transport, and splicing [82–84]. Lastly, RBPs are emerging as critical modulators of every hallmark of cancer and its druggable prediction through ML methods will provide a better understanding of potentially unveil new targets for cancer therapy and prognostic biomarkers.

## CONCLUSIONS

The current study proposed a new prediction model for druggable proteins by using as inputs the amino acid composition descriptors of protein sequences for 13 Machine Learning classifiers (with or without feature selection/dimension reduction of features). The best classifier was provided by SVM using only 200 selected tri-amino acid composition descriptors (TC). The high performance of the model is based on a mean AUROC of 0.975 ± 0.003 and a mean accuracy of 0.929 ± 0.006 (3-fold cross-validation). Thousands of protein sequences related to cancer have been used to predict the druggability. The top 10 BC OncoOmics essential proteins predicted as druggable were CDK4, AP1S1, POLE, HMMR, RPL5, PALB2, TIMP1, RPL22, NFKB1 and TOP2A; the top 10 cancer-driving proteins predicted as druggable were TLL2, FAM47C, SAGE1, HTR1E, MACC1, ZFR2, VMA21, DUSP9, CTNNA3 and GABRG1; and the top 10 RNA-binding proteins predicted as druggable were PLA2G1B, CPEB2, NOL6, LRRC47, CTTN, CORO1A, SCAF11, KCTD12, DDX43 and TMPO. Nevertheless, most of them do not currently have clinical trials. Finally, this powerful model predicts several druggable proteins which should be deeply studied to find better therapeutic targets and thus improve clinical trials.

## ABBREVIATIONS

FDA: US Food & Drug Administration;
IM: integral membrane;
SPTM: single pass transmembrane;
ML: machine learning;
BC: breast cancer;
RBPs: RNA-binding proteins;
AC: amino acid composition;
DC: di-amino acid composition;
TC: tri-amino acid composition;
GNB: Gaussian naive bayes;
SVM: support vector machine;
RBF: radial basis functions;
PCA: principal component analysis;
KNN: k-nearest neighbors algorithm;
LDA: linear discriminant analysis;
LG: logistic regression;
MLP: multilayer perceptron;
DT: decision tree;
RF: random forest;
GB: gradient boosting;
CV: cross-validation;
AUROC: area under the receiver operating characteristics;
SD: standard deviation;
FDR: false discovery rate;
GO: gene ontology;
PharmGKB: pharmacogenomics knowledgebase;
KEGG: Kyoto Encyclopedia of Genes and Genomes.

## ACKNOWLEDGMENTS

This work was supported by a) the Collaborative Project in Genomic Data Integration (CICLOGEN) PI17/01826 funded by the Carlos III Health Institute from the Spanish National plan for Scientific and Technical Research and Innovation 2013-2016 and the European Regional Development Funds (FEDER) - “A way to build Europe”; b) the General Directorate of Culture, Education and University Management of Xunta de Galicia ED431D 2017/16 and “Drug Discovery Galician Network” Ref. ED431G/01 and the “Galician Network for Colorectal Cancer Research” (Ref. ED431D 2017/23); c) the Spanish Ministry of Economy and Competitiveness for its support through the funding of the unique installation BIOCAI (UNLC08-1E-002, UNLC13-13-3503) and the European Regional Development Funds (FEDER) by the European Union; d) the Consolidation and Structuring of Competitive Research Units - Competitive Reference Groups (ED431C 2018/49), funded by the Ministry of Education, University and Vocational Training of the Xunta de Galicia endowed with EU FEDER funds; e) research grants from Ministry of Economy and Competitiveness, MINECO, Spain (FEDER CTQ2016-74881-P), Basque government (IT1045-16), and kind support of Ikerbasque, Basque Foundation for Science; and, f) Sociedad Latinoamericana de Farmacogenómica y Medicina Personalizada (SOLFAGEM).

## AUTHORS’ CONTRIBUTIONS

ALC, ACA and CRM conceived the subject, the conceptualization of the study and wrote the manuscript. ALC, ACA, CMCS and CRM did data curation and supplementary data. CRM and JD built the models using machine learning. JD, AP, HGD, CPyM, YPC and ET gave conceptual advice and valuable scientific input. JD, AP, HGD, CPyM, YPC, ET and CRM supervised the project. ALC and CPyM did funding acquisition. Finally, all authors reviewed the manuscript.

## AVAILABILITY OF DATA AND MATERIALS

All data generated during this study are included in this published article, and the scripts are available as a free repository at https://github.com/muntisa/machine-learning-for-druggable-proteins.

## REFERENCES

1. Venter JC, Smith HO, Adams MD (2015) The Sequence of the Human Genome. Clinical Chemistry 61:1207–1208

2. Barbarino JM, Whirl-Carrillo M, Altman RB, Klein TE (2018) PharmGKB: A worldwide resource for pharmacogenomic information. Wiley Interdiscip Rev Syst Biol Med 10:e1417

3. Whirl-Carrillo M, McDonagh EM, Hebert JM, et al (2012) Pharmacogenomics knowledge for personalized medicine. Clin Pharmacol Ther 92:414–417

4. Uhlén M, Fagerberg L, Hallström BM, et al (2015) Proteomics. Tissue-based map of the human proteome. Science 347:1260419

5. Kandoi G, Acencio ML, Lemke N (2015) Prediction of Druggable Proteins Using Machine Learning and Systems Biology: A Mini-Review. Front Physiol 6:366

6. Brown D, Superti-Furga G (2003) Rediscovering the sweet spot in drug discovery. Drug Discov Today 8:1067–1077

7. Cheng AC, Coleman RG, Smyth KT, et al (2007) Structure-based maximal affinity model predicts small-molecule druggability. Nat Biotechnol 25:71–75

8. Gashaw I, Ellinghaus P, Sommer A, Asadullah K (2012) What makes a good drug target? Drug Discov Today 17 Suppl:S24–30

9. Oprea TI, Bologa CG, Brunak S, et al (2018) Unexplored therapeutic opportunities in the human genome. Nat Rev Drug Discov 17:377

10. Wishart DS, Knox C, Guo AC, et al (2006) DrugBank: a comprehensive resource for in silico drug discovery and exploration. Nucleic Acids Res 34:D668–72

11. Wishart DS, Knox C, Guo AC, et al (2008) DrugBank: a knowledgebase for drugs, drug actions and drug targets. Nucleic Acids Res 36:D901–6

12. Costa PR, Acencio ML, Lemke N (2010) A machine learning approach for genome-wide prediction of morbid and druggable human genes based on systems-level data. BMC Genomics 11 Suppl 5:S9

13. Blanco JL, Porto-Pazos AB, Pazos A, Fernandez-Lozano C (2018) Prediction of high anti-angiogenic activity peptides in silico using a generalized linear model and feature selection. Sci Rep 8:15688

14. Wei L, Zhou C, Chen H, et al (2018) ACPred-FL: a sequence-based predictor using effective feature representation to improve the prediction of anti-cancer peptides. Bioinformatics

15. Concu R, M Natália D, Munteanu CR, González-Díaz H (2019) PTML Model of Enzyme Subclasses for Mining the Proteome of Biofuel Producing Microorganisms. Journal of Proteome Research 18:2735–2746

16. Martínez-Arzate SG, Tenorio-Borroto E, Pliego AB, et al (2017) PTML Model for Proteome Mining of B-Cell Epitopes and Theoretical–Experimental Study of Bm86 Protein Sequences from Colima, Mexico. Journal of Proteome Research 16:4093–4103

17. Fernandez-Lozano C, Cuiñas RF, Seoane JA, et al (2015) Classification of signaling proteins based on molecular star graph descriptors using Machine Learning models. J Theor Biol 384:50–58

18. Munteanu CR, Pedreira N, Dorado J, et al (2014) LECTINPred: web Server that Uses Complex Networks of Protein Structure for Prediction of Lectins with Potential Use as Cancer Biomarkers or in Parasite Vaccine Design. Molecular Informatics 33:276–285

19. Fernández-Blanco E, Aguiar-Pulido V, Munteanu CR, Dorado J (2013) Random Forest classification based on star graph topological indices for antioxidant proteins. Journal of Theoretical Biology 317:331–337

20. Zhu M, Gao L, Li X, et al (2009) The analysis of the drug-targets based on the topological properties in the human protein-protein interaction network. J Drug Target 17:524–532

21. Jeon J, Nim S, Teyra J, et al (2014) A systematic approach to identify novel cancer drug targets using machine learning, inhibitor design and high-throughput screening. Genome Med 6:57

22. Li Z-C, Zhong W-Q, Liu Z-Q, et al (2015) Large-scale identification of potential drug targets based on the topological features of human protein-protein interaction network. Anal Chim Acta 871:18–27

23. Laenen G, Thorrez L, Börnigen D, Moreau Y (2013) Finding the targets of a drug by integration of gene expression data with a protein interaction network. Mol Biosyst 9:1676–1685

24. Emig D, Ivliev A, Pustovalova O, et al (2013) Drug target prediction and repositioning using an integrated network-based approach. PLoS One 8:e60618

25. Yao L, Rzhetsky A (2008) Quantitative systems-level determinants of human genes targeted by successful drugs. Genome Res 18:206–213

26. Yildirim MA, Goh K-I, Cusick ME, et al (2007) Drug-target network. Nat Biotechnol 25:1119–1126

27. Cao D-S, -S. Cao D, Xiao N, et al (2015) Rcpi: R/Bioconductor package to generate various descriptors of proteins, compounds and their interactions. Bioinformatics 31:279–281

28. Hao J, Ho TK (2019) Machine Learning Made Easy: A Review of Scikit-learn Package in Python Programming Language. Journal of Educational and Behavioral Statistics 44:348–361

29. Brewka G (1996) Artificial intelligence—a modern approach by Stuart Russell and Peter Norvig, Prentice Hall. Series in Artificial Intelligence, Englewood Cliffs, NJ. The Knowledge Engineering Review 11:78–79

30. Cover T, Hart P (1967) Nearest neighbor pattern classification. IEEE Transactions on Information Theory 13:21–27

31. Cristianini N (2004) Fisher Discriminant Analysis (Linear Discriminant Analysis). Dictionary of Bioinformatics and Computational Biology

32. Patle A, Chouhan DS (2013) SVM kernel functions for classification. 2013 International Conference on Advances in Technology and Engineering (ICATE)

33. Peduzzi P, Concato J, Kemper E, et al (1996) A simulation study of the number of events per variable in logistic regression analysis. Journal of Clinical Epidemiology 49:1373–1379

34. Rosenblatt F (1961) PRINCIPLES OF NEURODYNAMICS. PERCEPTRONS AND THE THEORY OF BRAIN MECHANISMS

35. Swain PH, Hauska H (1977) The decision tree classifier: Design and potential. IEEE Transactions on Geoscience Electronics 15:142–147

36. Breiman L (2001) Machine Learning. 45:5–32

37. Chen T, Guestrin C (2016) XGBoost. Proceedings of the 22nd ACM SIGKDD International Conference on Knowledge Discovery and Data Mining - KDD ‘16

38. Friedman JH (2002) Stochastic gradient boosting. Computational Statistics & Data Analysis 38:367–378

39. Hughes G (1968) On the mean accuracy of statistical pattern recognizers. IEEE Transactions on Information Theory 14:55–63

40. Breiman L (1996) Bagging predictors. Machine Learning 24:123–140

41. Jolliffe IT (1986) Principal Component Analysis. Springer Series in Statistics

42. Law V, Knox C, Djoumbou Y, et al (2014) DrugBank 4.0: shedding new light on drug metabolism. Nucleic Acids Res 42:D1091–7

43. Tonks NK (2006) Protein tyrosine phosphatases: from genes, to function, to disease. Nature Reviews Molecular Cell Biology 7:833–846

44. López-Cortés A, Paz-y-Miño C, Guerrero S, et al OncoOmics approaches to reveal essential genes in breast cancer: a panoramic view from pathogenesis to precision medicine

45. Repana D, Nulsen J, Dressler L, et al (2019) The Network of Cancer Genes (NCG): a comprehensive catalogue of known and candidate cancer genes from cancer sequencing screens. Genome Biology 20

46. Hentze MW, Castello A, Schwarzl T, Preiss T (2018) A brave new world of RNA-binding proteins. Nat Rev Mol Cell Biol 19:327–341

47. Chawla NV, Bowyer KW, Hall LO, Kegelmeyer WP (2002) SMOTE: Synthetic Minority Over-sampling Technique. Journal of Artificial Intelligence Research 16:321–357

48. Bradley AP (1997) The use of the area under the ROC curve in the evaluation of machine learning algorithms. Pattern Recognition 30:1145–1159

49. Reimand J, Isserlin R, Voisin V, et al (2019) Pathway enrichment analysis and visualization of omics data using g:Profiler, GSEA, Cytoscape and EnrichmentMap. Nat Protoc 14:482–517

50. Carvalho-Silva D, Pierleoni A, Pignatelli M, et al (2019) Open Targets Platform: new developments and updates two years on. Nucleic Acids Res 47:D1056–D1065

51. Huang DW, Sherman BT, Lempicki RA (2009) Systematic and integrative analysis of large gene lists using DAVID bioinformatics resources. Nat Protoc 4:44–57

52. Ogata H, Goto S, Sato K, et al (1999) KEGG: Kyoto Encyclopedia of Genes and Genomes. Nucleic Acids Research 27:29–34

53. Slenter DN, Kutmon M, Hanspers K, et al (2018) WikiPathways: a multifaceted pathway database bridging metabolomics to other omics research. Nucleic Acids Res 46:D661–D667

54. Posey JE, Harel T, Liu P, et al (2017) Resolution of Disease Phenotypes Resulting from Multilocus Genomic Variation. N Engl J Med 376:21–31

55. Ivanov AA, Revennaugh B, Rusnak L, et al (2018) The OncoPPi Portal: an integrative resource to explore and prioritize protein-protein interactions for cancer target discovery. Bioinformatics 34:1183–1191

56. Li Z, Ivanov AA, Su R, et al (2017) The OncoPPi network of cancer-focused protein-protein interactions to inform biological insights and therapeutic strategies. Nat Commun 8:14356

57. Tsherniak A, Vazquez F, Montgomery PG, et al (2017) Defining a Cancer Dependency Map. Cell 170:564–576.e16

58. Uhlen M, Oksvold P, Fagerberg L, et al (2010) Towards a knowledge-based Human Protein Atlas. Nature Biotechnology 28:1248–1250

59. López-Cortés A, Paz-Y-Miño C, Cabrera-Andrade A, et al (2018) Gene prioritization, communality analysis, networking and metabolic integrated pathway to better understand breast cancer pathogenesis. Sci Rep 8:16679

60. Ding L, Bailey MH, Porta-Pardo E, et al (2018) Perspective on Oncogenic Processes at the End of the Beginning of Cancer Genomics. Cell 173:305–320.e10

61. Huang K-L, Mashl RJ, Wu Y, et al (2018) Pathogenic Germline Variants in 10,389 Adult Cancers. Cell 173:355–370.e14

62. Bailey MH, Tokheim C, Porta-Pardo E, et al (2018) Comprehensive Characterization of Cancer Driver Genes and Mutations. Cell 174:1034–1035

63. Liu J, Lichtenberg T, Hoadley KA, et al (2018) An Integrated TCGA Pan-Cancer Clinical Data Resource to Drive High-Quality Survival Outcome Analytics. Cell 173:400–416.e11

64. Hoadley KA, Yau C, Hinoue T, et al (2018) Cell-of-Origin Patterns Dominate the Molecular Classification of 10,000 Tumors from 33 Types of Cancer. Cell 173:291–304.e6

65. Tamborero D, Rubio-Perez C, Deu-Pons J, et al (2018) Cancer Genome Interpreter annotates the biological and clinical relevance of tumor alterations. Genome Med 10:25

66. Guerrero S, López-Cortés A, Indacochea A, et al (2018) Analysis of Racial/Ethnic Representation in Select Basic and Applied Cancer Research Studies. Sci Rep 8:13978

67. López-Cortés A, Guerrero S, Redal MA, et al (2017) State of Art of Cancer Pharmacogenomics in Latin American Populations. Int J Mol Sci 18.: https://doi.org/10.3390/ijms18060639

68. Quinones LA, Lavanderos MA, Cayun JP, et al (2014) Perception of the usefulness of drug/gene pairs and barriers for pharmacogenomics in Latin America. Curr Drug Metab 15:202–208

69. Paz-Y-Miño C, Robles P, Salazar C, et al (2016) Positive association of the androgen receptor CAG repeat length polymorphism with the risk of prostate cancer. Mol Med Rep 14:1791–1798

70. López-Cortés A, Echeverría C, Oña-Cisneros F, et al (2015) Breast cancer risk associated with gene expression and genotype polymorphisms of the folate-metabolizing MTHFR gene: a case-control study in a high altitude Ecuadorian mestizo population. Tumor Biology 36:6451–6461

71. López-Cortés A, Cabrera-Andrade A, Oña-Cisneros F, et al (2018) Breast Cancer Risk Associated with Genotype Polymorphisms of the Aurora Kinase a Gene (AURKA): a Case-Control Study in a High Altitude Ecuadorian Mestizo Population. Pathology & Oncology Research 24:457–465

72. López-Cortés A, Leone PE, Freire-Paspuel B, et al (2018) Mutational Analysis of Oncogenic AKT1 Gene Associated with Breast Cancer Risk in the High Altitude Ecuadorian Mestizo Population. BioMed Research International 2018:1–10

73. López-Cortés A, Jaramillo-Koupermann G, Muñoz MJ, et al (2013) Genetic polymorphisms in MTHFR (C677T, A1298C), MTR (A2756G) and MTRR (A66G) genes associated with pathological characteristics of prostate cancer in the Ecuadorian population. Am J Med Sci 346:447–454

74. Paz-y-Miño C, Muñoz MJ, López-Cortés A, et al (2010) Frequency of polymorphisms pro198leu in GPX-1 gene and ile58thr in MnSOD gene in the altitude Ecuadorian population with bladder cancer. Oncol Res 18:395–400

75. Harbeck N, Penault-Llorca F, Cortes J, et al (2019) Breast cancer. Nat Rev Dis Primers 5:66

76. López-Cortés A, Paz-y-Miño C, Guerrero S, et al (2019) Pharmacogenomics, biomarker network, and allele frequencies in colorectal cancer. The Pharmacogenomics Journal

77. García-Cárdenas JM, Guerrero S, López-Cortés A, et al (2019) Post-transcriptional Regulation of Colorectal Cancer: A Focus on RNA-Binding Proteins. Frontiers in Molecular Biosciences 6

78. Iadevaia V, Gerber AP (2015) Combinatorial Control of mRNA Fates by RNA-Binding Proteins and Non-Coding RNAs. Biomolecules 5:2207–2222

79. Wurth L, Gebauer F (2015) RNA-binding proteins, multifaceted translational regulators in cancer. Biochim Biophys Acta 1849:881–886

80. Keene JD (2007) RNA regulons: coordination of post-transcriptional events. Nat Rev Genet 8:533–543

81. Wurth L (2012) Versatility of RNA-Binding Proteins in Cancer. Comp Funct Genomics 2012:178525

82. Burd CG, Dreyfuss G (1994) Conserved structures and diversity of functions of RNA-binding proteins. Science 265:615–621

83. Lukong KE, Chang K-W, Khandjian EW, Richard S (2008) RNA-binding proteins in human genetic disease. Trends in Genetics 24:416–425

84. Kechavarzi B, Janga SC (2014) Dissecting the expression landscape of RNA-binding proteins in human cancers. Genome Biol 15:R14

